# Transfer learning predicts species-specific drug interactions in emerging pathogens

**DOI:** 10.1101/2024.06.04.597386

**Authors:** Carolina H. Chung, David C. Chang, Nicole M. Rhoads, Madeline R. Shay, Karthik Srinivasan, Mercy A. Okezue, Ashlee D. Brunaugh, Sriram Chandrasekaran

## Abstract

Machine learning (ML) algorithms are necessary to efficiently identify potent drug combinations within a large candidate space to combat drug resistance. However, existing ML approaches cannot be applied to emerging and under-studied pathogens with limited training data. To address this, we developed a transfer learning and crowdsourcing framework (TACTIC) to train ML models on data from multiple bacteria. TACTIC was built using 2,965 drug interactions from 12 bacterial strains and outperformed traditional ML models in predicting drug interaction outcomes for species that lack training data. Top TACTIC model features revealed genetic and metabolic factors that influence cross-species and species-specific drug interaction outcomes. Upon analyzing ∼600,000 predicted drug interactions across 9 metabolic environments and 18 bacterial strains, we identified a small set of drug interactions that are selectively synergistic against Gram-negative (e.g., *A. baumannii*) and non-tuberculous mycobacteria (NTM) pathogens. We experimentally validated synergistic drug combinations containing clarithromycin, ampicillin, and mecillinam against *M. abscessus*, an emerging pathogen with growing levels of antibiotic resistance. Lastly, we leveraged TACTIC to propose selectively synergistic drug combinations to treat bacterial eye infections (endophthalmitis).

## Introduction

Antibiotic resistance is becoming a global health crisis as growing resistance outpaces current antibiotic drug development. In the United States alone, more than 2.8 million antibiotic-resistant infections occur each year, with more than 35,000 cases resulting in death^1^. These infections are caused by a diverse set of bacterial pathogens that pose major health threats, especially if left unchecked. Some of the most concerning species include carbapenem-resistant *Acinetobacter baumannii*, multidrug-resistant Pseudomonas aeruginosa, and drug-resistant *Mycobacterium tuberculosis* (henceforth *M. tb*)^1,2^. At the same time, no new classes of antibiotics have been brought to market for decades^3^.

One possible solution for overcoming antibiotic resistance is to design synergistic drug combination therapies^4^. Such combinations could engage multiple cellular targets to suppress growing resistance, which is difficult to achieve with a single active compound. A major drawback, however, is the challenge in exploring a vast combinatorial space that exponentially increases when considering new drugs or dosage levels. Due to the empirical nature of designing drug combination therapies, the discovery of new combined regimens has been slow. Most notably, the four-drug regimen course that is standard for treating tuberculosis (TB) has not changed in 50 years, which has led to growing resistance^5^. There is therefore a dire need for alternative approaches that can efficiently screen and prioritize promising drug combinations.

Machine learning (ML) has recently been leveraged to design synergistic drug combinations for various complex diseases^6–14^. In context of antibiotic resistance, an approach called INDIGO was developed to find combination therapies effective against Escherichia coli and *M. tb*^15,16^. Specifically, omics data for *E. coli* and *M. tb* treated with individual drugs, along with corresponding drug combination assay data, were used to construct two separate models predictive of drug interaction outcomes (e.g., synergy) for each organism. The *E. coli* INDIGO model was further extended to predict drug interaction outcomes for Staphylococcus aureus by leveraging omics data for genes that are orthologous between the two species. Although INDIGO was shown to yield predictions that correlate with experimental data, the INDIGO models are not highly predictive for species with limited data compared to *E. coli* (e.g. *Mycobacteria*).

Within this study, we leverage transfer learning and crowdsourcing to construct predictive models of drug interactions for emerging clinically relevant pathogens, for which little or no data exists for training traditional ML models. Transfer learning is a ML concept where a model trained on one task is re-applied to solve a different but similar task, typically after some finetuning^17^. Crowdsourcing describes the process of amalgamating input from various information sources to complete a common task^18^. We integrate these concepts onto a modeling foundation inspired by INDIGO to develop a new approach called TACTIC: <u>T</u>ransfer learning <u>A</u>nd <u>C</u>rowdsourcing to predict <u>T</u>herapeutic Interactions <u>C</u>ross- species. We show that TACTIC generates more accurate predictions than INDIGO for 12 phylogenetically diverse bacterial strains. We then interpret the TACTIC model to explain the genetic and metabolic drivers of cross-species and species-specific drug interaction outcomes. Finally, we apply TACTIC to predict drug combinations that show narrow- spectrum synergy against multiple groups of pathogenic bacteria and antagonism against common commensal microbes.

## Results

### Drug interaction data collection

To build and validate our framework, we compiled drug interaction data from 17 publications^15,16,19–33^ (**Figure 1**, **Table 1**). In total, we curated 2965 drug interactions involving 88 conditions (86 drugs and 2 media) measured across 12 strains representative of six species: *Acinetobacter baumannii, E. coli, M. tb, Pseudomonas aeruginosa, S. aureus, and Salmonella enterica Serovar Typhimurium* (henceforth *S. Typhimurium*) (**Figure 1**, **Data S1 and S2**). Drug interactions were quantified based on the Loewe Additivity^34^ or Bliss Independence^35^ model. Based on the drug interaction classification scheme defined for each study, our pooled data consists of 913 synergistic (31%), 826 neutral (28%), and 1226 antagonistic (41%) interactions (**Figure *1***). Our pooled data is representative of two- to ten-way drug combinations, although more than 85% (N = 2522) consists of pairwise interactions.

**Figure 1.**
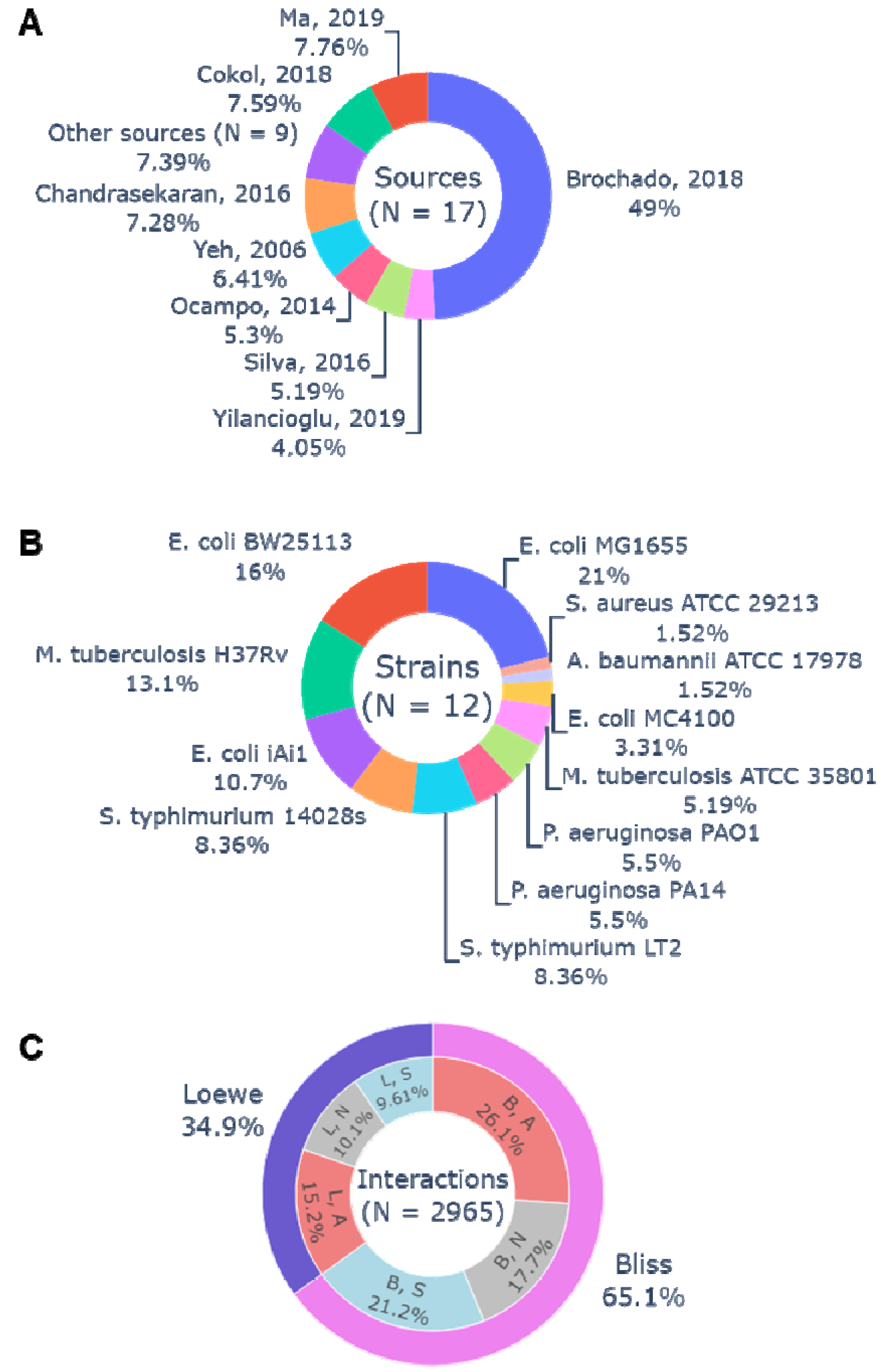
Drug interaction data collection for TACTIC. (**A**) Drug interaction data collected from 17 published sources was used to build TACTIC models. (**B**) This data collection measured outcomes in 12 bacterial strains representative of six organisms: Acinetobacter baumannii, Escherichia coli, Mycobacterium tuberculosis, Pseudomonas aeruginosa, Salmonella enterica serovar Typhimurium, and Staphylococcus aureus. (**C**) Drug interaction outcomes were measured using the Loewe Additivity and/or the Bliss Independence model, and data distribution was found to be relatively balanced between the three outcome classes: synergy (S), neutral (N), and antagonism (A).

**Table 1.** TACTIC data collection. Drug interaction outcome measurements gathered from 17 independent studies. N_total_ = total number of interactions, N_high_ = number of high-order interactions, Ref.: reference.

| Organism | Strain | $N_{\text{total}}$ | $N_{\text{high}}$ | Model | Interaction Outcome | | | | | | Ref. |
| --- | --- | --- | --- | --- | --- | --- | --- | --- | --- | --- | --- |
|  |  |  |  |  | *Synergy* |  | *Neutral* |  | *Antagonism* |  |  |
| *A. baumannii* | ATCC 17978 | 45 | 30 | Loewe | 23 | (0.51) | 10 | (0.22) | 12 | (0.27) | <sup>30</sup> |
| *E. coli* | BW25113 | 473 | - | Both | 117 | (0.25) | 155 | (0.33) | 201 | (0.42) | <sup>27,33</sup> |
|  | MC4100 | 98 | 26 | Both | 12 | (0.12) | 24 | (0.25) | 62 | (0.63) | <sup>21</sup> |
|  | MG1655 | 623 | 185 | Both | 123 | (0.20) | 218 | (0.35) | 282 | (0.45) | <sup>15,23,30,32</sup> |
|  | iAi1 | 316 | - | Bliss | 116 | (0.37) | 66 | (0.21) | 134 | (0.42) | <sup>27</sup> |
| *M. tuberculosis* | ATCC 35801 | 154 | 134 | Bliss | 85 | (0.55) | 41 | (0.27) | 28 | (0.18) | <sup>22</sup> |
|  | H37Rv | 389 | 68 | Loewe | 181 | (0.47) | 56 | (0.14) | 152 | (0.39) | <sup>16,19,20,24–26,28,29,31</sup> |
| *P. aeruginosa* | PAO1 | 163 | - | Bliss | 44 | (0.27) | 47 | (0.29) | 72 | (0.44) | <sup>27</sup> |
|  | PA14 | 163 | - | Bliss | 51 | (0.31) | 54 | (0.33) | 58 | (0.36) | <sup>27</sup> |
| *S. aureus* | ATCC 29213 | 45 | - | Loewe | 5 | (0.11) | 23 | (0.51) | 17 | (0.38) | <sup>15</sup> |
| *S. Typhimurium* | 14028s | 248 | - | Bliss | 77 | (0.31) | 68 | (0.27) | 103 | (0.42) | <sup>27</sup> |
|  | LT2 | 248 | - | Bliss | 79 | (0.32) | 64 | (0.26) | 105 | (0.42) | <sup>27</sup> |

Out of the 1370 unique drug combinations represented in our data collection, 757 (∼55%) were measured more than once (**Figure S1A**, **Data S3**). For drug combinations repeatedly measured in different strains (N = 664), the vast majority (>86%) either agreed on a single outcome class (N = 215) or disagreed between neighboring class labels (synergy vs. neutral or neutral vs. antagonism, N = 357) (**Figure S1B**). For example, ciprofloxacin (CIP) combined with spectinomycin (SPE) was found to have antagonistic activity across seven different strains. CIP combined with chloramphenicol (CHL) was also found to be antagonistic or neutral across seven strains. Polymyxin B (PMB) paired with benzalkonium (BZK), rifampicin (RIF), or verapamil (VER) was found to be synergistic across six strains. These drug interactions may thus show broad-spectrum synergistic activity, which may be a strategic therapeutic option to treat infections where the causative agent is unknown.

In contrast, only ∼14% (N = 92) of drug interactions measured in more than one organism were found to have diverging outcomes across different strains. Of these, 45 drug combinations were measured in a single study that quantified interactions using the Bliss Independence model across up to six strains representing *E. coli*, *P. aeruginosa*, and *S. typhimurium*^27^. Some of these interactions generally displayed similar interaction outcomes across most strains except one. For example, A22 combined with mitomycin C (MMC) was antagonistic or additive across most strains except *E. coli* BW25113 (**Figure S1C**). Another example is MMC combined with RIF, which was antagonistic across most strains except S. Typhimurium 14028s, for which it was synergistic.

Interestingly, some drug interactions were highly specific to the strain even within the same species. For example, BZK combined with EGCG was synergistic against *E. coli* BW25113 yet antagonistic against *E. coli* iAi1 (**Figure S1C**). Similarly, MMC combined with procaine (PRC) was synergistic against *E. coli* BW25113 yet antagonistic against *E. coli* iAi1. This combination was also found to be synergistic against *P. aeruginosa* PAO1 yet antagonistic against *P. aeruginosa* PA14. These drug interactions may represent those with narrow-spectrum activity, which could be leveraged to treat bacterial infections where the causative agent is known while minimizing unintended damage to other bacteria (e.g., the host microbiome).

### Benchmarking TACTIC against INDIGO

TACTIC is an extension of INDIGO^15^, a computational approach for predicting drug interaction outcomes between antibacterial agents. Our approach specifically expands on the orthology aspect, which enables the application of a dataset measured for one organism (e.g., *E. coli*) to another (e.g., *S. aureus*). Unlike INDIGO models, which are trained on data for a single organism (i.e., *E. coli* or *M. tb*), a TACTIC model is trained on omic and drug interaction data measured in multiple organisms (**Figure 2**). When modeling multiple strains using TACTIC, each organism is represented by its unique list of genes and orthologs. The ability to incorporate multi-species drug interaction and omics data enables the model to learn from a larger and more varied set of data compared to when training on data from a single organism. This allows TACTIC to perform transfer learning and infer drug interaction outcomes for a new strain of interest based on gene orthology information alone. All other steps involved in developing a predictive model (e.g., defining ML model features, the ML algorithm used) were preserved from the original INDIGO model and are reprised in the Methods section.

**Figure 2.**
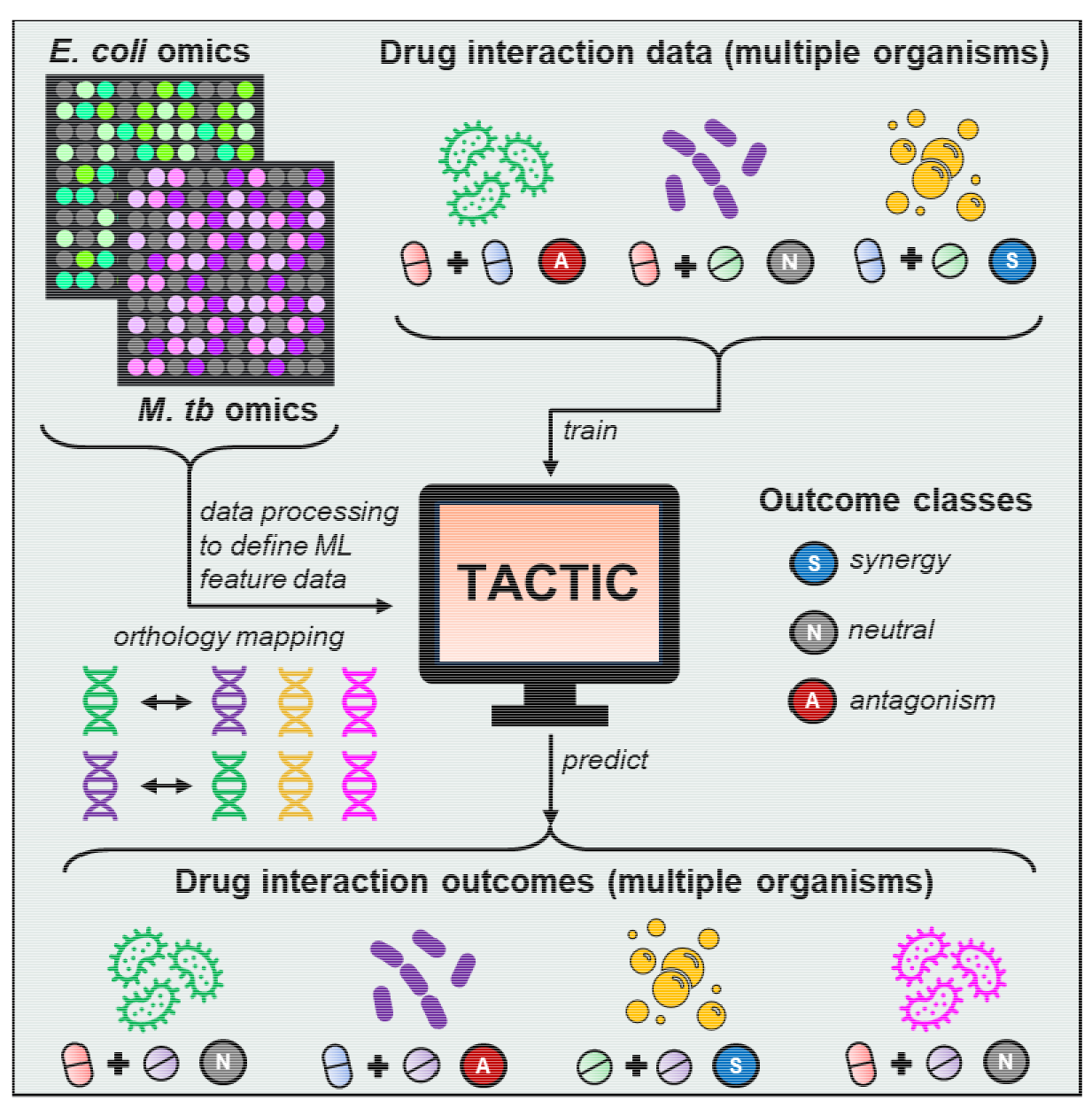
TACTIC approach schematic. The *<u>T</u>ransfer learning <u>A</u>nd <u>C</u>rowdsourcing to predict <u>T</u>herapeutic Interactions <u>C</u>ross-species* (TACTIC) approach uses single drug response and drug interaction data to train a machine learning (ML) model that predicts drug interaction outcomes for multiple organisms. Single drug response data is processed to define ML features for a given set of drug combinations. Orthology mapping allows the transfer of drug response data measured in one organism to another by conserving data between orthologous genes.

We benchmarked TACTIC against INDIGO by evaluating their accuracy in predicting drug interaction outcomes for each of the 12 bacterial strains for which data was collected above. For INDIGO, we directly used the previously constructed *E. coli*^15^ and *M. tb*^16^ models to generate predictions for all 2965 drug interactions. For the TACTIC approach, we trained one model for each strain-specific dataset, where drug interactions measured for the strain of interest (N) were set aside for testing while the remaining drug interaction data (2965-N) was used for training a model. Model accuracy was measured based on the Spearman rank correlation and the mean squared error (MSE) between model predictions and experimentally measured drug interaction scores.

We found that TACTIC models were considerably better at predicting drug interaction outcomes for A. baumannii, *P. aeruginosa*, S. Typhimurium, and most *E. coli* strains compared to INDIGO models (**Figure 3**). Unsurprisingly, the INDIGO models built using *E. coli* and *M. tb* data yielded slightly more accurate predictions than TACTIC for *E. coli* MC4100 and *M. tb*, respectively. Interestingly, all three models were less accurate in predicting drug interaction outcomes for *S. aureus* (Spearman R < 0.4 and MSE > 1 for all models). This could be because *S. aureus* (Gram-positive) is phylogenetically distant from all other bacteria considered within this benchmarking analysis (Gram-negative or Mycobacterium strains). In other words, transfer learning across all models is limited by the small overlap of orthologous genes between *S. aureus*, *E. coli*, and *M. tb*.

**Figure 3.**
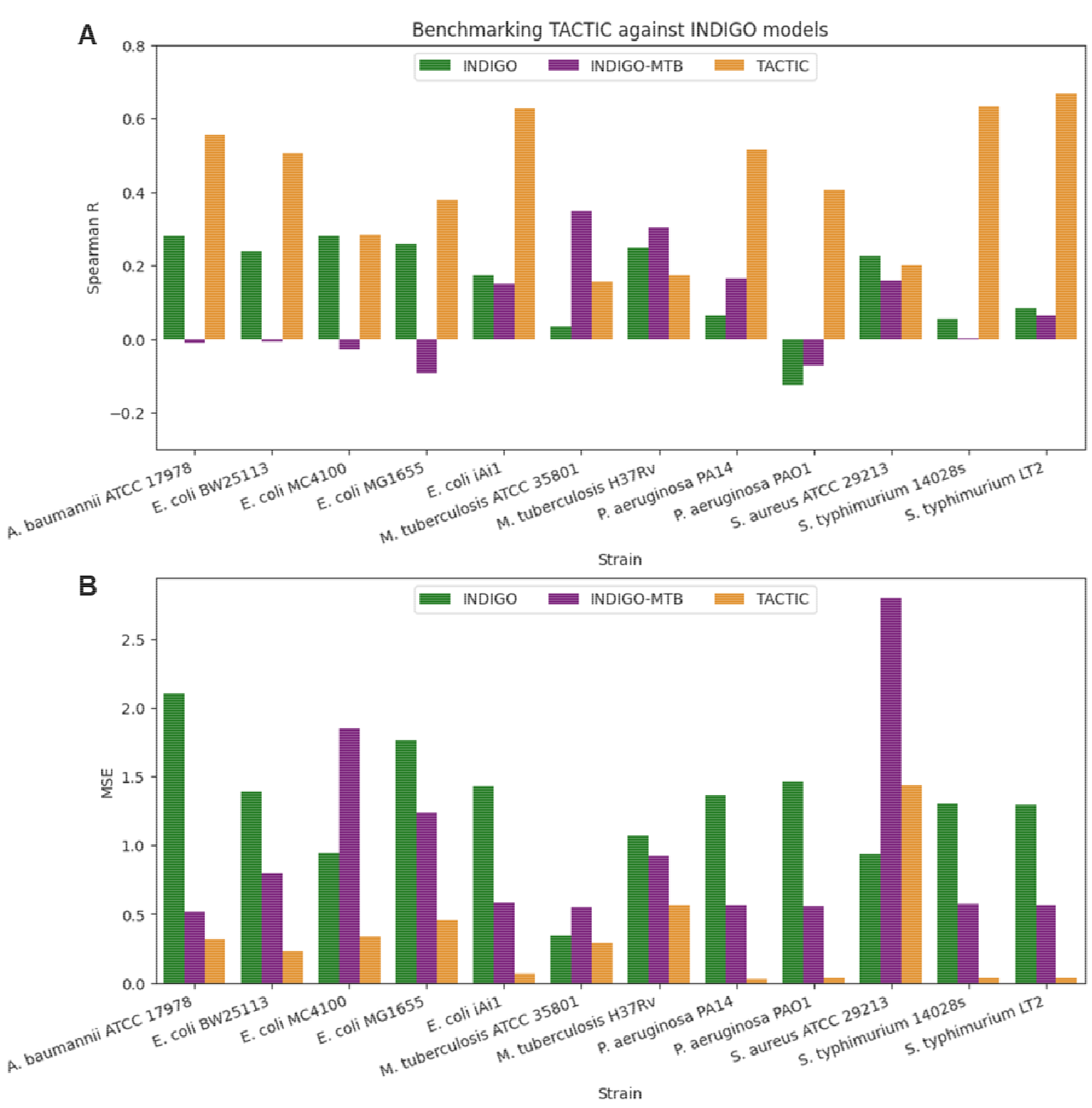
Benchmarking TACTIC against INDIGO. Strain-specific drug interaction outcome prediction accuracy was compared between TACTIC models and INDIGO models developed from *E. coli* or *M. tb* data^15,16^. Model performance was quantified based on (**A**) the Spearman rank correlation and (**B**) the mean squared error (MSE) between model predictions and experimentally measured drug interaction scores.

### Genetic predictors of cross-species and species-specific drug interaction outcomes

Having confirmed that TACTIC provides an improvement over INDIGO in predicting drug interaction outcomes cross-species, we next trained a TACTIC model with the entire data collection on the 2965 drug interactions measured across 12 bacterial strains. Using this fully trained TACTIC model, we sought to uncover which features the model most relied on for generating predictions and how changes in these feature values explained synergistic or antagonistic outcomes. For this task, we first determined the set of ML features, ranked by decreasing importance score, that explained 95% of the variance in model predictions (see Methods for details). We then conducted a gene set enrichment analysis of all genes associated with the top ML features (N ∼ 1400) against the KEGG database^36^, which revealed four pathways that are significantly enriched (**Figure S2**, hypergeometric test, adjusted p-value < 0.05). Of note, all pathways relate to bacterial metabolism, with the biosynthesis of amino acids enriched by genes specific to both *E. coli* MG1655 and *M. tb* H37Rv.

To gain a more fine-grained insight, we evaluated how the values for ML features associated with genes belonging to these pathways varied between synergistic and antagonistic interactions (see Methods for details). We determined 11 genes in central metabolic pathways for which feature values significantly differed between synergistic and antagonistic interactions (**Table *2***, **Figure S3**). The trends for all 11 genes indicate that drug interactions involving at least one drug that perturbs central metabolic pathways (i.e., glycolysis, TCA cycle and oxidative phosphorylation) likely result in antagonistic outcomes. These findings may align with a longstanding hypothesis that fully intact and active bacterial metabolism may potentiate drug efficacy^37,38^; therefore, gene perturbations that attenuate metabolic activity may decrease combined antibiotic potency and lead to antagonistic interactions.

**Table 2.** *E. coli* MG1655 gene knockouts associated with antagonistic outcomes. KO: knockout.

| Locus | Gene | Description | Primary Pathway | KO Fitness |
| --- | --- | --- | --- | --- |
| b0115 | aceF | pyruvate dehydrogenase, E2 subunit | Glycolysis / Gluconeogenesis | Positive |
| b0118 | acnB | hypothetical protein | Citrate cycle (TCA cycle) |  |
| b1136 | icd | isocitrate dehydrogenase |  |  |
| b3737 | atpE | ATP synthase Fo complex subunit c | Oxidative phosphorylation |  |
| b4042 | dgkA | diacylglycerol kinase | Glycerolipid metabolism |  |
| b3041 | ribB | 3,4-dihydroxy-2-butanone-4-phosphate synthase | Riboflavin metabolism | Positive |
| b0844 | ybjI | 5-amino-6-(5-phospho-D-ribitylamino) uracil phosphatase |  | Negative |
| b3620 | waaF | ADP-heptose--LPS heptosyltransferase 2 | Lipopolysaccharide biosynthesis | Positive |
| b3625 | waaY | lipopolysaccharide core heptose (II) kinase |  | Negative |
| b3628 | waaB | UDP-D-galactose: (glucosyl)lipopolysaccharide-1,6-D-galactosyltransferase |  |  |
| b3632 | waaQ | lipopolysaccharide core heptosyltransferase 3 |  |  |

We next investigated associations between ML features and species-specific drug interaction outcomes by analyzing 32 drug combinations measured in 6 different strains within the same study^27^. Within this subset, we found that the deviation in cross-species drug interaction outcomes aligns with the phylogenetic distance for combinations, especially for those involving bacteriostatic-bacteriostatic interactions (**Figure S4, Data S2**). This may indicate that bacteriostatic drug effects are more dependent on the genetic state of a given organism, as opposed to bactericidal agents which are believed to be more dependent on the phenotypic (e.g., metabolic) state of a bacterial cell^37,39–41^. This hypothesis aligns with our reanalysis of experimental measurements of the impact of various carbon sources on drug interactions^42^. We observed a larger variation in drug interactions for the combination of cefoxitin (bactericidal) paired with tobramycin (bactericidal) compared to cefoxitin paired with tetracycline (bacteriostatic) on *E. coli* growth across 95 carbon sources (**Figure S5**). However, a larger investigation of combined drug efficacy based on the mode of action (bacteriostatic vs. bactericidal) across defined metabolic conditions is necessary to confirm this trend.

### Interaction landscape of ∼3600 drug pairs in 18 strains across 9 media conditions

We next applied the fully trained TACTIC model to generate drug interaction predictions across new strains and media conditions. Since metabolic genes were top features in our model, we sought to understand the degree to which drug interaction outcomes can differ across distinct organisms and growth environments. We included all 12 strains for which the TACTIC model was trained on as well as two non-tuberculous mycobacteria (NTM) species (*Mycobacterium abscessus* and *Mycobacterium smegmatis*) and four bacteria representative of gut microbiome commensals (*Bacteroides vugatus, Eubacterium eigens, Eubacterium rectale, and Lactobacillus rhammosus*). These new strains were selected based on growing concerns over NTM infections^43^ and studies that have deduced the species-level identities of commensal gut microbes^44,45^, for which there is increasing appreciation over their symbiotic relationship with human hosts^46–49^. For the media conditions, we accounted for 9 nutrient sources that were represented in the input data for TACTIC. These included seven carbon sources (acetate, glucosamine, glucose, glycerol, maltose, N-acetylglucosamine, succinate), one nitrogen source (ammonium chloride), and one nutrient-rich condition (LB).

In total, we generated ∼600,000 outcome predictions spanning all possible drug pairs among 86 drugs (N = 3655) across 18 bacterial strains and 9 media conditions (**Data S4**). Given the large scale of this data, we applied principal component analysis (PCA) to inspect how each strain contextualized within each media condition cluster among one another (**Figure 4**). Interestingly, we found that differences in predicted outcomes were primarily driven by the media context, where predictions made in certain media conditions (e.g., N-acetylglucosamine) clustered within a narrow range across the first principal component (PC1). The patterns seen across PC1 indicate that the growth environment may sometimes have a larger impact on the combined drug effect across a diverse range of bacteria. Nonetheless, we observed that clustering patterns along the second principal component (PC2) were strongly driven by the strain identity, with some organisms clustering along both PC1 and PC2 (e.g., *S. aureus*, Mycobacteria). To further understand strain-specific differences in predicted drug interaction outcomes, we decided to conduct strain-to-strain comparisons for predictions contextualized in LB, the media condition where strain-to-strain variance was the highest (**Table S1**, see Methods for details).

**Figure 4.**
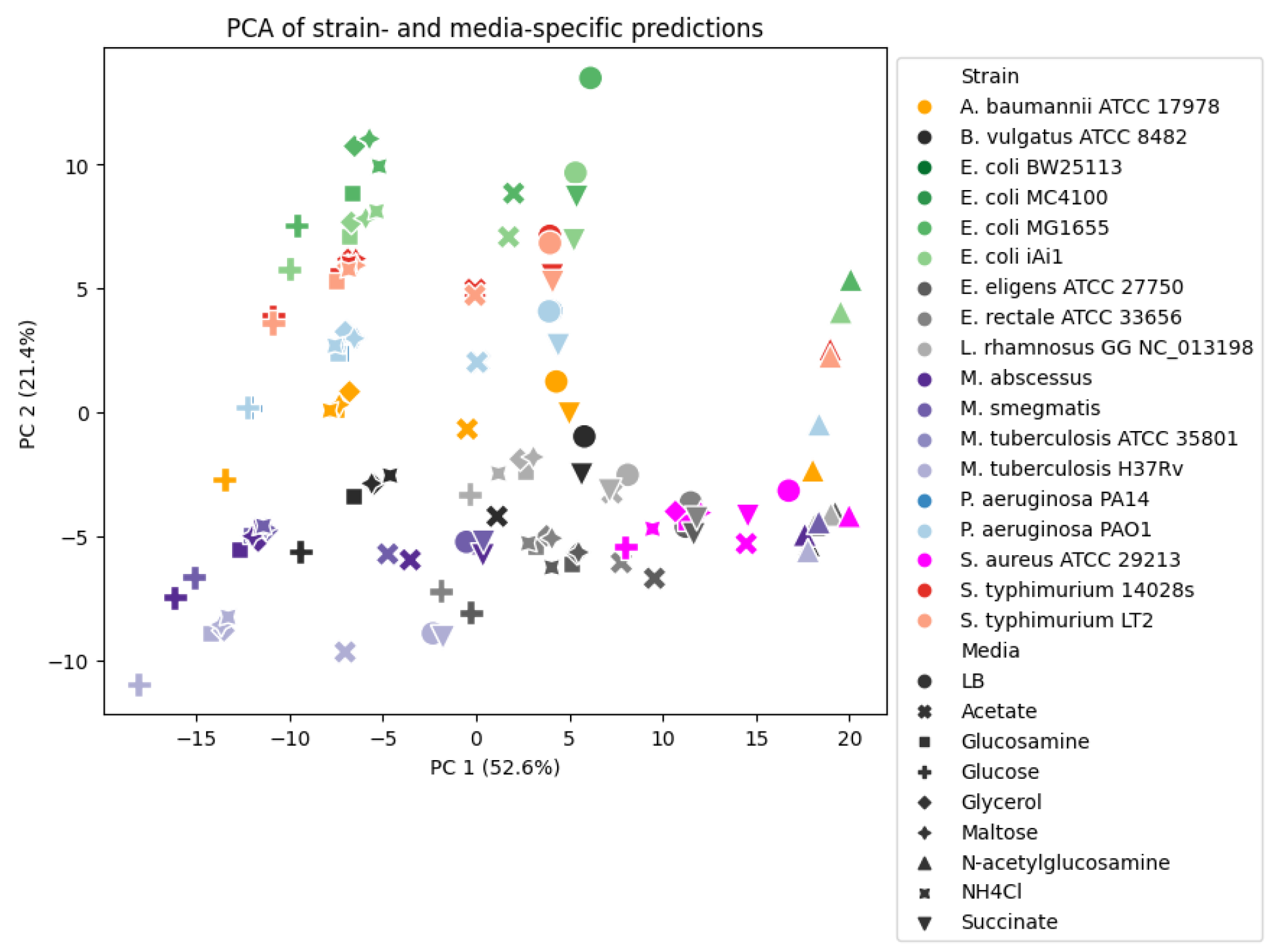
Principal component analysis (PCA) of TACTIC prediction landscape. PCA of pairwise drug interaction predictions specific to 18 bacterial strains (differentiated by marker colors) contextualized within 9 media conditions (differentiated by marker shape) reveals clustering patterns driven by media- (PC1) and strain- (PC2) specific factors. NH4Cl: ammonium chloride.

Using TACTIC predictions on 3655 drug interaction outcomes across 18 bacterial strains contextualized in LB, we first evaluated the similarity between strain-to-strain predictions. By defining similarity as the Pearson correlation between strain-specific predictions, we found that drug interaction outcome similarities are primarily based on the Gram stain of an organism (**Figure 5**). Of note, outcome predictions for *S. aureus* were the most dissimilar from all other strain-specific predictions (lowest correlation < 0.6). Unsurprisingly, strain-specific predictions for the same species were most similar to one another, and Mycobacteria-specific predictions clustered amongst each other. Of note, strain-specific predictions seemed to cluster based on the strain-specific relationship with the host (i.e., pathogen vs. commensal) within Gram stain groups. This finding supported the potential of identifying drug combinations with selective synergy against pathogenic bacteria within our strain-specific prediction landscape.

**Figure 5.**
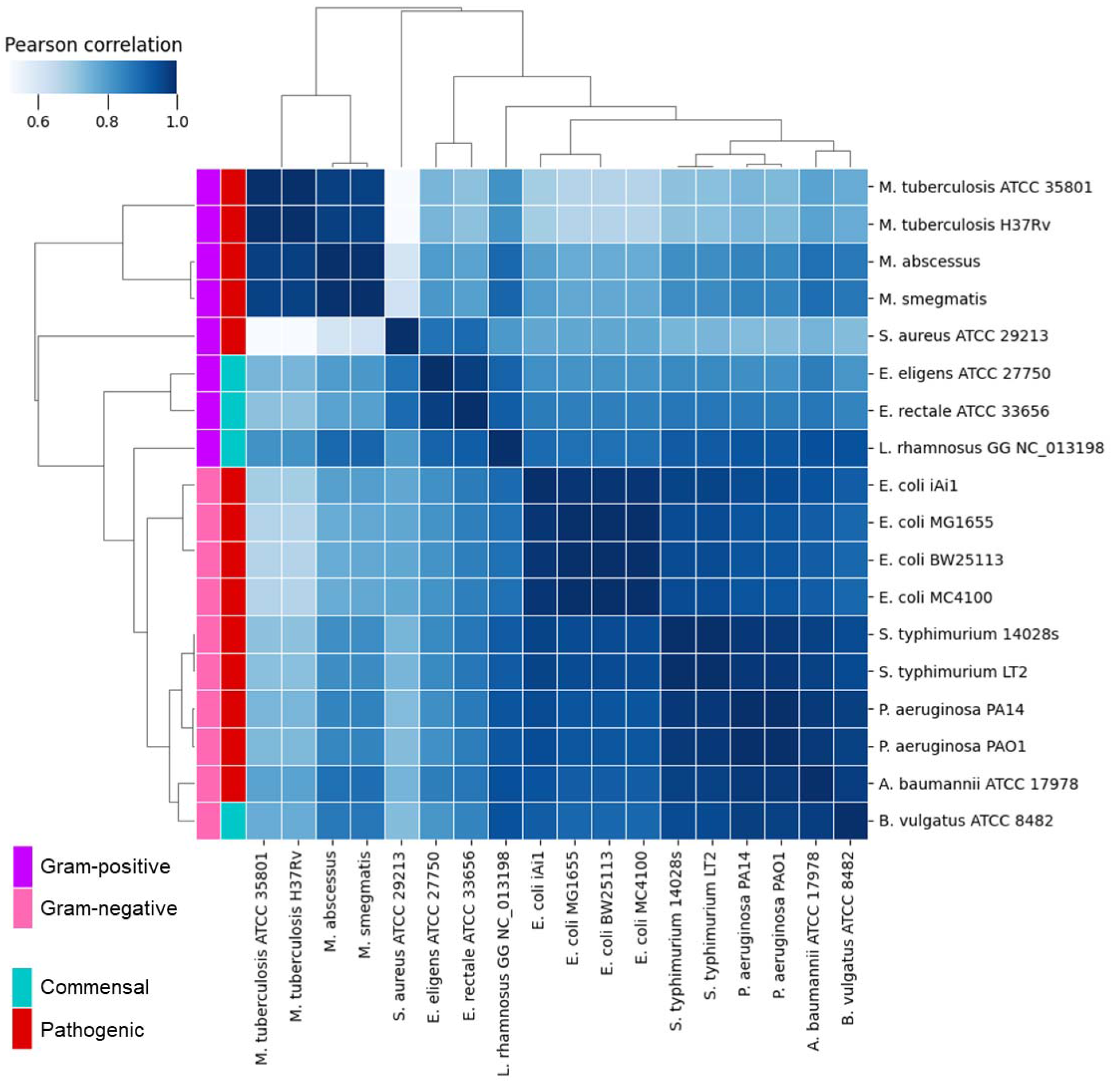
Correlation heatmap for strain-specific TACTIC model predictions in LB. Drug interaction outcome predictions for all possible drug pairs between 86 drugs (N = 3,655) were generated across 18 bacterial strains by the full TACTIC model. The Pearson correlation for strain-to-strain predictions was calculated for all strain pairs and visualized as a clustered heatmap.

### Identifying narrow-spectrum drug synergies against distinct pathogen groups

Prior to scanning the LB prediction landscape for narrow-spectrum synergies, we first evaluated how TACTIC predictions aligned with existing evidence on drug interaction outcomes measured in organisms accounted for in our model^50–69^. We found that TACTIC correctly predicted broad-spectrum synergy for most drug combinations determined to be synergistic across a wide range of bacteria, regardless of the combined mechanism of action (**Table 3**). The only exceptions were ampicillin combined with gentamicin (AMP– GEN) and polymyxin B combined with trimethoprim (PMB–TMP), which TACTIC predicted to have selective synergy against specific bacterial groups.

**Table 3.** TACTIC prediction agreement with literature-based evidence. Comparison between 10 drug interaction outcomes predicted by TACTIC against consensus derived from prior studies. Refer to **Data S2** for more information on abbreviated drug names.

| Interaction | Mechanism | Organism Measured | Literature Consensus | TACTIC Prediction |
| --- | --- | --- | --- | --- |
| AMP-AMK | Beta lactam and aminoglycoside | Gram-negative bacteria<sup>50,51,62</sup><br>*S. aureus*<sup>50</sup> | Broad-spectrum synergy | Broad-spectrum synergy |
| AMK-AMX |  | *E. coli*<sup>63</sup> |  |  |
| AMK-AZT |  | *P. aeruginosa*<sup>63–66</sup> |  |  |
| AZT-TOB |  | *P. aeruginosa*<sup>67</sup> |  |  |
| AMP-GEN |  | Gram-negative bacteria<sup>68</sup> | Selective synergy in Gram-negative bacteria | Selective synergy in *E. coli* and *S. Typhimurium* |
| AMP-AZT | Beta lactam pair | Gram-negative bacteria<sup>52,69</sup> | Selective synergy in Gram-negative bacteria | Broad-spectrum synergy |
| SMM-TMP | Double targeting of folic acid biosynthesis | Gram-positive and Gram-negative bacteria<sup>53</sup> | Broad-spectrum synergy | Broad-spectrum synergy |
| CLA-RIF | Target RNA and protein biosynthesis | NTM species<sup>54–57</sup> | Selective synergy in Mycobacteria | Selective synergy in Mycobacteria |
| PMB-RIF | Target RNA and lipopolysaccharide biosynthesis | Gram-negative bacteria<sup>58</sup> | Selective synergy in Gram-negative bacteria | Selective synergy in Gram-negative and Mycobacteria |
| PMB-TMP | Target lipopolysaccharide and folic acid biosynthesis | Gram-positive and Gram-negative bacteria<sup>59–61</sup> | Broad-spectrum synergy | Selective synergy in Mycobacteria and *S. aureus* |

We also found that TACTIC predictions generally aligned with literature evidence pointing to selective synergistic drug combinations. For example, TACTIC correctly predicts that clarithromycin combined with rifampicin (CLA–RIF) is synergistic against Mycobacteria^54–57^. Interestingly, TACTIC predicts that polymyxin B combined with rifampicin (PMB–RIF) is selectively synergistic against Gram-negative and Mycobacteria species; though prior evidence of synergy against Gram-negative bacteria exists^58^, the effect of this drug combination has yet to be measured in Mycobacteria. Another interesting drug combination is ampicillin paired with another beta-lactam drug aztreonam (AMP–AZT), which was previously shown to be synergistic against various Gram-negative pathogens^52,69^; yet TACTIC predicts broad-spectrum synergy for this combination, which warrants its evaluation against other bacteria.

To identify drug combinations with narrow-spectrum synergy, we compared TACTIC model predictions for commensals vs. three strain groups: all pathogens, Gram-negative pathogens, and NTM pathogens. For these comparisons, we imposed two criteria to define narrow-spectrum synergy. The first was to select drug combinations that are predicted to be additive or antagonistic (predicted interaction score (IS) >= 0) across all commensal strains. The second criterium selected pathogen-specific synergies based on one of four thresholds: strong comprehensive synergy (predicted IS < -0.2 for all strains), comprehensive synergy (predicted IS < 0 for all strains), strong average synergy (mean predicted IS < -0.2 for all strains), and average synergy (mean predicted IS < 0 for all strains).

For the first comparison involving all pathogens, only the average synergy threshold returned narrow-spectrum synergies (**Figure 6**). A visual inspection of the top 20 narrow-spectrum synergies revealed that broad synergistic outcomes are hard to achieve for all pathogens, most likely because this group includes both Gram-negative and Gram-positive species. For the second comparison with Gram-negative pathogens, 34 drug combinations were predicted to have comprehensive narrow-spectrum synergy (**Figure 6**), with stronger synergies predicted against *E. coli* and *S. Typhimurium* strains compared to A. baumannii or *P. aeruginosa* strains. For the third comparison, 16 drug combinations were predicted to have strong comprehensive narrow-spectrum synergy against NTM pathogens (**Figure 6**). Interestingly, most of these combinations (N = 14) include clarithromycin, which is known to have potent effect against NTM strains^70^.

**Figure 6.**
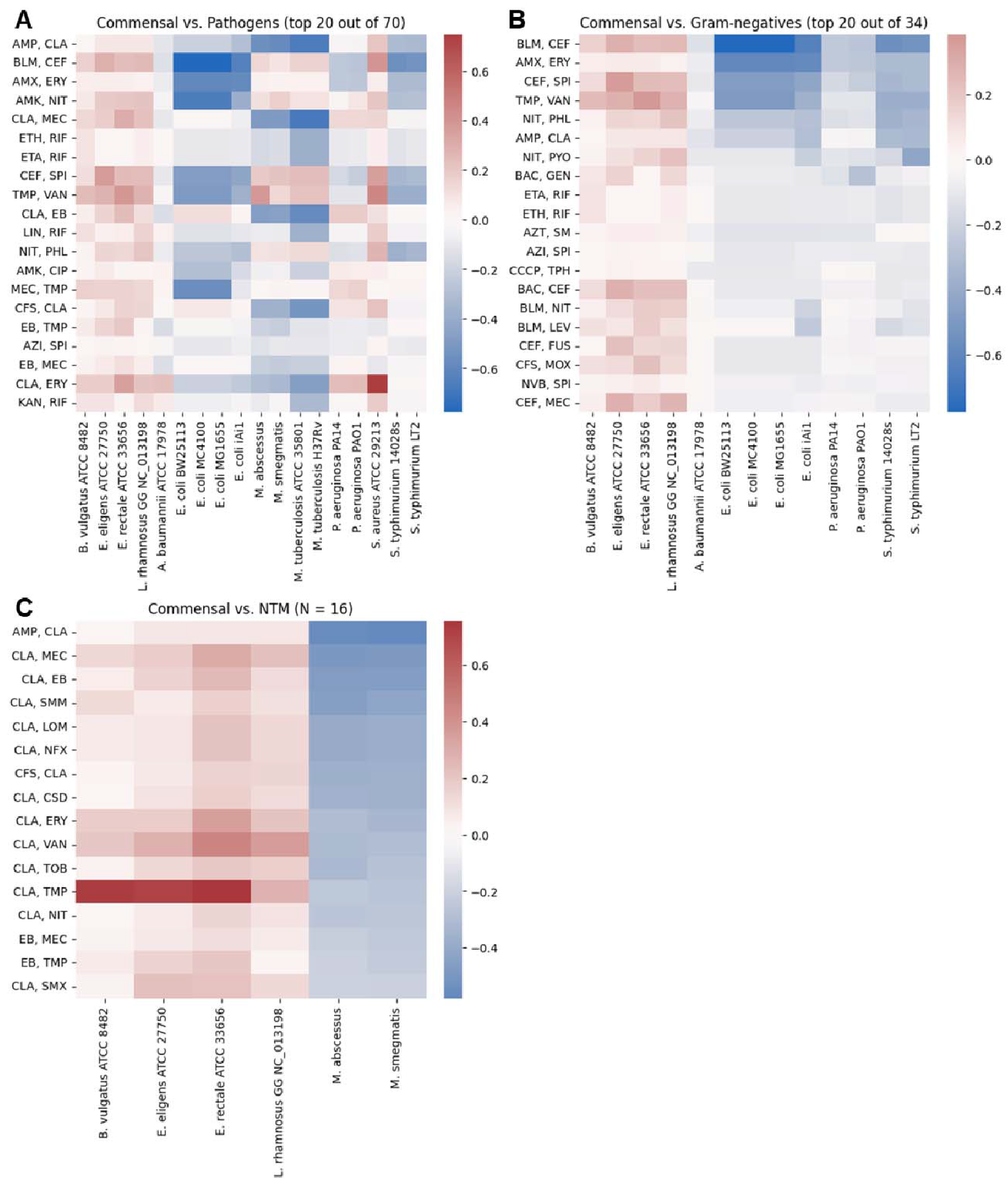
Narrow-spectrum synergy predictions against pathogenic bacteria. Narrow-spectrum synergy was defined by additive or antagonistic predictions (predicted IS >= 0) across all commensal strains and synergistic predictions (predicted IS < 0) across pathogenic strains. In total, (**A**) 70 drug pairs are predicted to have general synergy (mean IS < 0) against all pathogenic strains, (**B**) 34 drug pairs are predicted to have comprehensive synergy (all IS < 0) against Gram-negative pathogens, and (**C**) 16 drug pairs are predicted to have strong comprehensive synergy (all IS < -0.2) against non-tuberculous mycobacteria (NTM) pathogens. The top 2 combinations (AMP-CLA, CLA-MEC) were chosen for experimental validation.

Given the lack of comprehensive drug interaction datasets for NTM, it is challenging to build ML models tailored for these pathogens. Considering that infections caused by NTM pathogens are on the rise^71^, with some species (e.g., *M. abscessus*) already developing concerning levels of antibiotic resistance^72^, we decided to experimentally confirm whether any of the 16 drug combinations identified by TACTIC were indeed synergistic against NTM species. For this investigation, we evaluated the effect of the top two synergistic predictions (AMP–CLA and CLA–MEC) against *M. abscessus* str. ATCC 19977 as well as two clinical isolates from a patient with cystic fibrosis (see Methods). Of note, the two clinical isolates exhibited differences in morphotypes (smooth vs. rough), which is known to influence pathogenesis and antibiotic resistance in *M. abscessus*^73^.

We confirmed that AMP–CLA was synergistic against *M. abscessus* ATCC 19977 and the smooth morphotype isolate but antagonistic in the rough morphotype isolate (**Figure 7**), suggesting that biophysical factors (i.e. morphology) may also influence drug interactions. We were unable to measure the combined effect for CLA–MEC due to an indeterminate minimal inhibitory concentration (MIC) for mecillinam (MEC). Hence, we measured the combined effect of all three antibiotics (AMP–CLA–MEC), where CLA and MEC were pre-combined in a 1:1 solution. Of note, TACTIC predicted that this three-drug combination also exhibits narrow-spectrum synergy against NTM pathogens (**Data S4**). Our experimental results show that AMP–CLA–MEC is synergistic against all three *M. abscessus* strains at high sub-inhibitory concentrations (**Figure 7**). In contrast, combinations of clarithromycin with beta-lactam drugs were previously found to be antagonistic or neutral in *E. coli, S. aureus*, and *Streptococcus pneumoniae*^15,74^. TACTIC also predicts that such combinations are weakly synergistic or antagonistic in non-Mycobacteria strains (**Figure S6**). Altogether, the experimental data agree with TACTIC predictions for the top selective synergies against *M. abscessus*.

**Figure 7.**
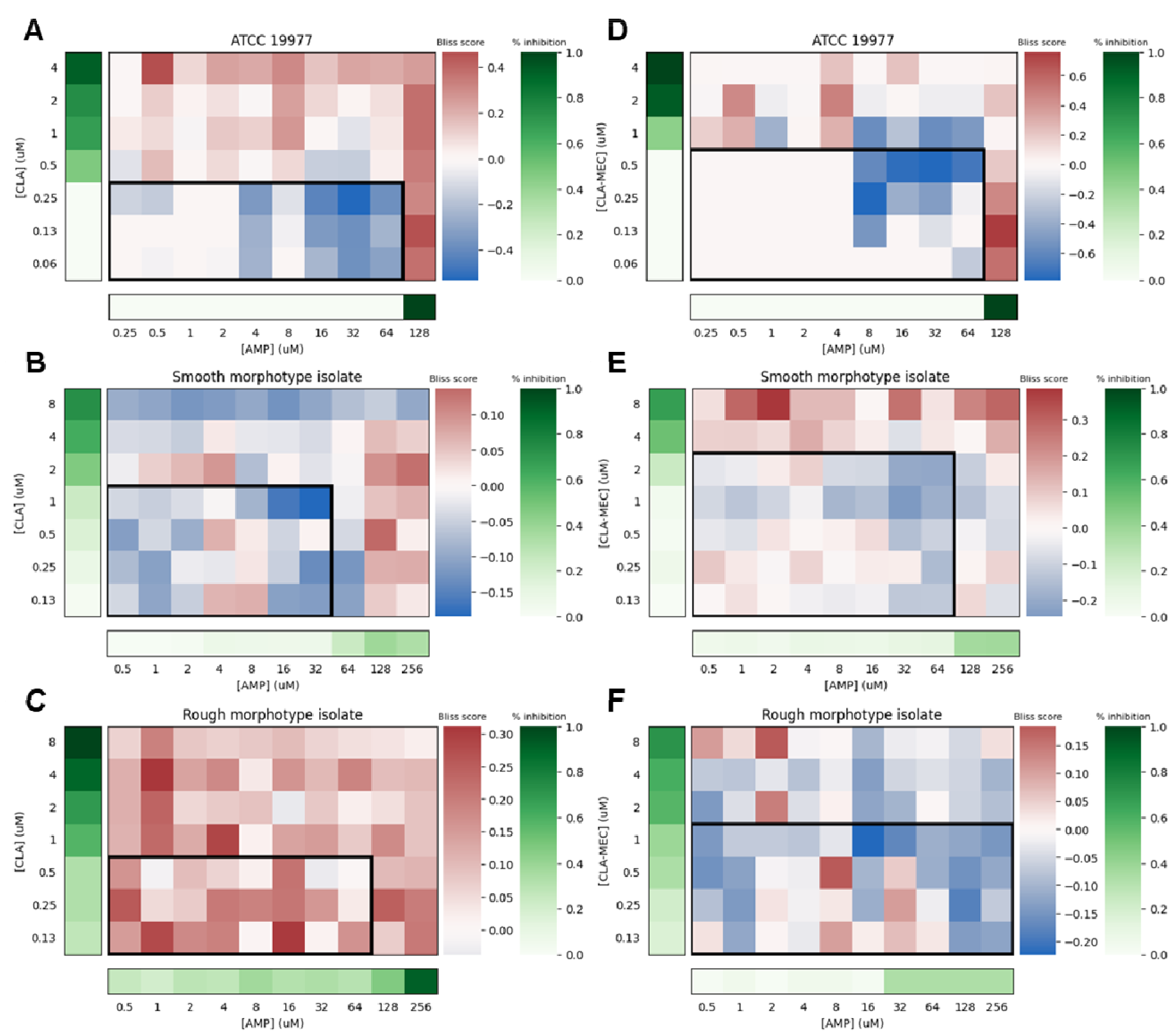
Experimental validation of synergy predictions for *M. abscessus*. (**A-C**) The combined effect of ampicillin paired with clarithromycin (AMP-CLA) was measured in *M. abscessus* ATCC 19977 and two clinical isolates with smooth or rough morphotypes. (**D-F**) The three-way combination between ampicillin, clarithromycin, and mecillinam (AMP-CLA-MEC) was also measured in the same three *M. abscessus* strains. For all checkerboards, the percentage of growth inhibition is shown for individual drug treatment wells while the Bliss score is reported for combination wells (see Methods for Bliss score calculation). Regions with subinhibitory levels (< 50%) of the drugs are highlighted with a black outline.

Motivated by the capacity to pinpoint synergistic drug combinations for the emerging pathogen *M. abscessus*, we next applied TACTIC to predict effective combination therapies against bacterial cases of endophthalmitis, a serious eye infection that can lead to blindness if improperly treated^75^. In addition to *P. aeruginosa*, the following species are considered key pathogens: Pseudomonas mendocina, Pseudomonas putida, Pseudomonas stutzeri, Neisseria spp., and Gemella haemolysans. To treat endophthalmitis, 15 antibiotics are typically used to clear bacterial infections (**Table S2**), with the combination of gatifloxacin and moxifloxacin (GAT–MOX) used as the first-line treatment. According to literature evidence on the ocular microbiome^76,77^, species belonging to the Corynebacterium, Propionibacterium, and Staphylococcus genera are predominantly found on the eye surface of healthy individuals.

Based on the combined information above, we applied TACTIC to predict drug interaction outcomes for all possible pairwise combinations between 73 drugs against 12 bacterial strains (**Table S3**). The drug selection included 12 out of the 15 clinically-used antibiotics for treating endophthalmitis that are present in the TACTIC model. In total, we generated more than 30,000 predictions that account for 2628 unique drug pairs across 12 strains (**Data S5**). GAT–MOX, which is commonly used as a broad-spectrum regimen to treat bacterial cases of endophthalmitis^78^, is predicted to be mildly synergistic against all *P. aeruginosa* and Neisseria strains, but antagonistic for the other Pseudomonas species. For the commensal strains (*C. kroppenstedtii* and *P. propionicum*), this combination is predicted to be additive (i.e., neutral). These predictions indicate that GAT–MOX may be an effective first-line therapy for cases of endophthalmitis primarily caused by *P. aeruginosa* or Neisseria spp. but not for other cases. Hence, GAT–MOX may not be the optimal first-line treatment due to its incomplete and weak coverage against all identified pathogenic strains.

Prior to scanning the TACTIC prediction landscape for alternative treatment options, we compared predictions against experimental data from studies that measured the efficacy of antibiotic combinations against clinical and drug-resistant strains of *P. aeruginosa*^79,80^ (**Figure S7**). Out of the 13 drug combinations involving polymyxin B (PMB), the TACTIC model correctly predicts that PMB yields stronger synergies when paired with aztreonam (AZT), fosfomycin (FOS), minocycline (MIN), and rifampicin (RIF) compared to other drugs. TACTIC model predictions also indicate that ceftazidime (CFZ) paired with tobramycin (TOB) is relatively more synergistic than CFZ combined with ciprofloxacin (CIP) or CIP–TOB, aligning with literature-based trends. Assured that TACTIC predictions align with existing evidence measured in *P. aeruginosa*, we identified 48 out of 2628 possible drug combinations having selective synergy against pathogenic bacteria (**Figure 8**, **Data S6**). Inspection of the ten most synergistic drug interactions against pathogenic strains revealed combinations that involve gentamicin (GEN) and CFZ, which are commonly used to treat endophthalmitis and are more effective against Gram-negative pathogens^81^. Interestingly, the remaining top synergistic set includes combinations involving at least one antibiotic active against the bacterial cell wall (mechanism of CFZ activity) but these combinations have not yet been investigated in treating bacterial endophthalmitis.

**Figure 8.**
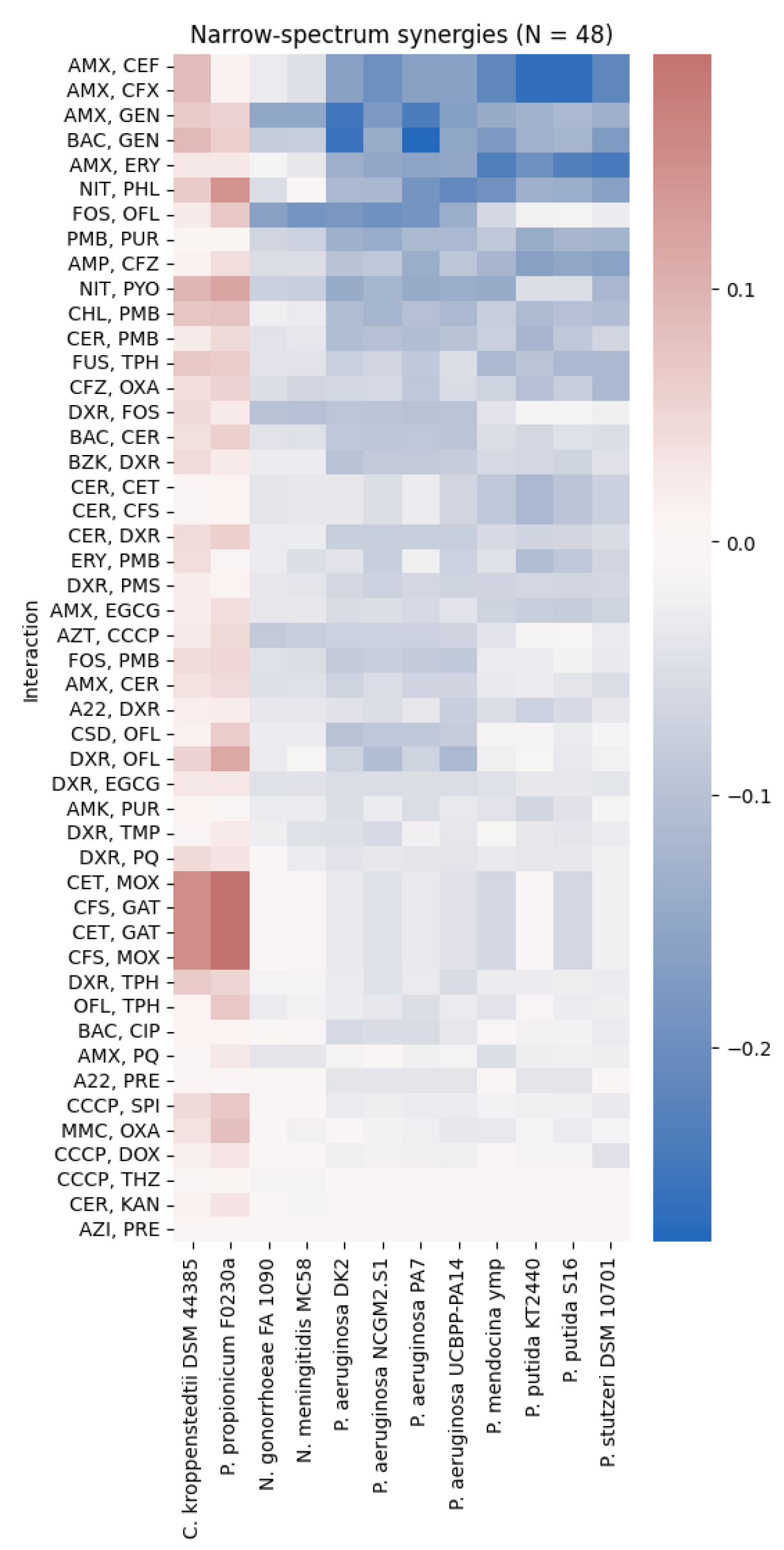
Drug combinations predicted to have narrow-spectrum synergy against bacterial agents of endophthalmitis. Out of 2,628 drug combinations, 48 were predicted to have narrow-spectrum synergy against causative agents for endophthalmitis (predicted IS < 0 against pathogenic strains, predicted IS ≥ 0 for commensal strains). The first two strains in the heatmap (C. kroppenstedtii and P. propionicum) are commensals. Refer to **Data S2** for more information on abbreviated drug names.

## Discussion

Here we demonstrate how data from model organisms can be leveraged for drug discovery in emerging pathogens. We introduce TACTIC, a new computational approach that leverages transfer learning and crowdsourcing via orthology mapping to learn drug-response patterns from diverse model organisms and predict strain-specific drug interaction outcomes in unseen strains. Using a data collection of ∼3000 drug interactions measured across a phylogenetically diverse set of bacterial strains (**Data S1**), we show that TACTIC models generate more accurate predictions cross-species compared to INDIGO^15^. The conservation of genes predictive of drug interactions across species enables accurate transfer learning with relatively less training data than a traditional single species model. In contrast, existing drug discovery models are primarily built using chemical structures, but this data cannot differentiate between pathogen strains, and models need to be rebuilt for each organism. TACTIC is also effective because it can observe more drug combinations and their interaction outcomes in various bacterial pathogens possessing differing levels of antimicrobial susceptibilities.

Based on a fully trained TACTIC model, we show that species-specific drug interaction outcomes are influenced by the phenotypic (e.g., metabolic state), genetic (e.g., Gram-stain), and pathogenic nature of a given organism. TACTIC identified several genes in amino acid metabolism to be predictive of drug interactions across species. Drug interactions in a wide range of organisms are influenced by their metabolic state, and while central metabolism is highly conserved, bacteria show distinct species-specific patterns of amino acid consumption^82^. Our results suggest that in addition to the causative pathogen strain, the in vivo metabolic environment should be considered while selecting antibiotic combination therapies.

We then use the TACTIC model to generate a comprehensive prediction landscape of interaction outcomes for ∼3600 drug pairs across 18 bacterial strains and 9 media conditions. By comparing TACTIC model predictions for commensal strains against different groups of pathogenic bacteria, we identified a small set of drug interactions that are predicted to have narrow-spectrum synergy against specific pathogens. Importantly, we experimentally validated top predictions for selective synergy against NTM pathogens (ampicillin–clarithromycin and ampicillin–clarithromycin–mecillinam) to be synergistic against *M. abscessus* strains. We further applied the TACTIC model to determine promising combinations with narrow-spectrum synergy for treating bacterial cases of endophthalmitis. Although these predictions have yet to be validated experimentally, they provide a manageable starting point for discovering combination therapies optimized to clear bacterial infections while minimizing detrimental effects on the commensal microflora of a human host.

The TACTIC approach provides a novel method to investigate strain-specific drug interaction outcomes, thus being the first computational framework that explores the existence of combination therapies with broad- or narrow-spectrum synergy. Broad-spectrum synergistic combinations are promising candidates for empirical antibiotic therapy when the causative pathogen is unknown. Although broad-spectrum antibiotic treatments are commonly used today and have high translational potential, they may increase antibiotic resistance and potentially damage the native microbiota, leading to a range of adverse effects^83,84^. There is hence a need for narrow-spectrum treatments, which are more effective once the causative pathogen is known. However, we underscore the fact that the prediction accuracy for TACTIC models may decrease for species that are phylogenetically distant (e.g., *S. aureus*) from organisms that are accounted for during model training (e.g., *E. coli* and *M. tb*). To address this limitation, omics data measuring the drug response in other Gram-positive organisms could be integrated into TACTIC models to improve prediction accuracy and extend application to a diverse range of bacterial species.

## Materials and Methods

### Experimental design

TACTIC predictions for combinations involving ampicillin sodium (Cayman Chemical Company, USA), clarithromycin (Cayman Chemical Company, USA), and mecillinam (Cayman Chemical Company, USA) were confirmed using a microbroth dilution checkerboard assay, which was performed according to CLSI standards^85^. Resazurin dye was used as an indicator of bacterial growth, where metabolism of the dye produces a color change from blue (absorbance maximum of 570 nm) to pink (absorbance maximum of 600 nm). *M. abscessus* ATCC 19977 was thawed from frozen stock, streaked onto Middlebrook 7H10 agar plate, and incubated for 72 hours at 37°C. Morphotype-matched (i.e., rough and smooth) clinical isolates were obtained from the University of Michigan Health System from the sputum of a de-identified patient with cystic fibrosis.

Antibiotics dissolved in Middlebrook 7H9 broth were incubated in combination with ATCC 19977, smooth morphotype, or rough morphotype inoculum (6x10^4^ CFU; determined using a standard curve) in a 96-well plate for 24 hours at 37°C and 150 RPM, at which point the resazurin dye was added. The absorbance of each well was read at 570 nm and 600 nm 24 hours following the addition of the dye. Each microplate additionally included a sterile control (broth and resazurin only) and a growth control (inoculum, broth and resazurin only). The percentage inhibition for each antibiotic combination was calculated by comparing the absorbance of the treated wells to that of the control wells. At least three separate replicates were averaged to obtain the final experimental results. The clinical isolates exhibited a baseline clarithromycin MIC of > 8 µg/mL and ampicillin MIC of > 264 µg/mL, as reported by the University of Michigan Health System. The ampicillin and clarithromycin combination exhibited inhibitory effects that were enhanced relative to the individual drugs, with the greatest effect observed in the smooth morphotype.

Experimentally measured drug interactions were quantified based on the Bliss Independence model^35^. This framework quantifies drug synergy based on the combined drug effect on cell growth by comparing the observed growth inhibition against the expected value, which is computed for a drug pair as follows:

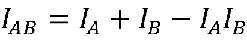

where I_AB_ is the expected growth inhibition achieved by the combination based on the inhibition achieved by each drug independently (I_A_ and I_B_). The Bliss score reported in **Figure 7** was defined as the difference between the expected and observed I_AB_, where negative scores indicate synergy.

### Data acquisition and curation

Two distinct data types were used to construct TACTIC models: drug response and drug interaction data. Drug response information was comprised of *E. coli* chemogenomic data^86^, where fitness of single-gene knockout strains was measured in response to individual stressors, and *M. tb* transcriptomic data^16^, which measured differential gene expression in response to individual stressors. Each omic dataset was previously z-score normalized within its originating study; hence, the normalized omic data was directly used for defining TACTIC model features. The drug interaction data comprised of 2965 combinations collected from 17 literature sources covering 12 bacterial strains (**Table *1***, **Data S1**). Drug combinations ranged between two- to ten-way interactions and were quantified based on the Loewe Additivity^34^ and/or the Bliss Independence^35^ model based on the originating study. A total of 178 unique drug interactions were measured based on both mathematical models, and the Spearman rank correlation between Bliss- and Loewe- based scores indicated a strong agreement between metrics (R = 0.466, p-value < 5e-11, **Figure S8**). This suggested that Bliss and Loewe drug interaction scores are concordant and could be trained together. Of note, the drug interaction data collection used in this study represents the subset for which the full set of ML features could be determined based on the omics datasets from which drug response information was extracted. For exploratory analysis of the data (**Figure *1* and S1**), each drug interaction was labeled into one of the three outcome classes (i.e., synergy, neutral, antagonism) based on the classification criteria specified in its originating publication. Continuous interaction score values were used for regression-based ML construction.

### Defining ML features for drug combinations

The method for defining ML features for a given drug combination is fully described in the INDIGO study^15^. Briefly, drug response data is first binarized to indicate substantial gene-level changes (i.e., z > 2 and z < -2) in cell response to a given stressor. The binarized vectors for all drugs involved in a given combination are then combined to define sigma and delta features for each gene, which mathematically represent the combined drug and drug-unique effect, respectively. Of note, a TACTIC model defines drug combination features based on two omic datasets that measure *E. coli* and *M. tb* drug response. To remove duplicated information captured within orthologous genes between these two organisms, a feature vector was defined by the sigma-delta transformation of the full set of *M. tb* genes and the sigma-delta transformation for genes unique to *E. coli*.

### Orthology mapping for Transfer learning

The *E. coli* and *M. tb* omics datasets were directly used to define ML features for drug combinations measured against *E. coli* str. K-12 substr. MG1655 (henceforth *E. coli* MG1655) and *M. tb* H37Rv. For all other organisms, ML features were defined by first determining the sigma-delta feature vector for a given drug combination then filtering this vector based on the orthology mapping between *E. coli* MG1655, *M. tb* H37Rv, and the given strain. Gene orthology was determined by mapping the *E. coli* MG1655 and *M. tb* H37Rv genomes against the genomes for all other strains based on genome annotation available in OrtholugeDB^87^, a database that infers orthologs based on the reciprocal-best-BLAST hit (RBBH) rate.

For all strain-specific predictions, the strain selection was based on the genomes that were available in OrtholugeDB for both pathogenic and commensal bacteria. For the endophthalmitis predictions, the 12 strains represent the following eight species: *P. aeruginosa*, *P. mendocina, P. putida, P. stutzeri, Neisseria gonorrhoeae, Neisseria meningitidis, Corynebacterium kroppenstedtii*, and *Propionibacterium propionicum* (**Table S3**). Of note, no genomes were available for species belonging to the Gemella genus, and the Staphylococcus genus was not included in the commensal set due to evidence of Staphylococci commonly being the causative agent for Gram-positive endophthalmitis cases^88^.

### TACTIC model construction using Random Forests

All TACTIC models were constructed in Python v3.10.8 (Python Software Foundation) using the regression-based Random Forests (RF) algorithm supplied through the scikit-learn package v1.2.0^89^. Of note, the RF algorithm was confirmed to yield the best performing TACTIC models compared to other regression-based ML algorithms, namely: linear regression, ridge regression, support vector regression, decision tree regression, and gradient boosting regression (**Figure S9**). Briefly, RF is an ensemble method comprised of multiple decision trees that independently learn to associate feature information (i.e., sigma-delta features for a given drug combination) to a target variable (i.e., the measured drug IS) during the training phase. Each decision tree is then tasked with estimating the target variable (i.e., drug interaction outcome) given feature information alone during the testing phase, and the average estimated value (i.e., the predicted drug IS) is returned by the RF algorithm. Randomization is factored into the model construction process via random sampling with replacement of the training dataset for each decision tree. This yields a diverse collection of decision trees that together reduce the variance between model predictions against true values for the target variable. Of note, all TACTIC models were constructed using the default parameter values for the RandomForestRegressor method in scikit-learn.

### TACTIC model interpretation via feature importance

Feature importance for the full TACTIC model was determined by using the built-in computation for the scikit-learn RF algorithm, which defines importance as the mean decrease in variance for a regression task. Based on this definition, the importance score for each feature is calculated by first constructing a model without the given feature then comparing its prediction accuracy against that of the model trained on all features. Specifically, importance is quantified as the difference in the mean squared error between model predictions and true values for the target variable. TACTIC model interpretation was completed by using the set of features ranked by decreasing importance score with a cumulative sum of 0.95, or the set of the most important features that explain 95% of the mean decrease in variance. In total, ∼2500 features associated with 1258 *E. coli* MG1655 genes and 411 *M. tb* H37Rv genes (both sets including 251 orthologous genes between the two organisms) met this criterium (**Data S7**).

Based on the KEGG pathways that were significantly enriched by the *E. coli* MG1655 and *M. tb* H37Rv genes identified above (**Figure S2**), we evaluated how the values for TACTIC model features that are associated with genes belonging to these pathways varied between synergistic and antagonistic interactions. Specifically, we conducted a two-sample Student’s t-test with equal variance and the chi-square test with Yate’s correction to compare sigma and delta feature value differences, respectively. Using a stringent significance level (adjusted p-value < 1E-6) based on the adjusted p-value distribution for all feature-specific tests (**Figure S10**), we determined 79 sigma or delta features that differed based on the drug interaction outcome type (**Data S8**). From this set, we identified 11 genes for which both sigma and delta feature values significantly different between synergistic and antagonistic interactions along the same direction for gene-specific changes (**Figure S3**).

### Finding associations between ML feature patterns and drug interaction outcomes

We assessed interaction scores for 32 drug combinations measured in all strains that were considered within the same study^27^. We defined strain-specific drug impact based on the equation below:

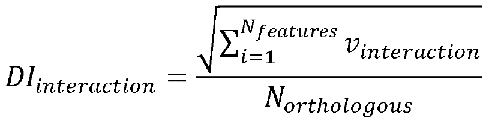

where DI_interaction_ is the drug impact for a given drug interaction, v_interaction_ is the vector of ML features determined for the given drug interaction, N_features_ is the total number of ML features, and N_orthologous_ is the number of *E. coli* MG1655 and *M. tb* H37Rv genes that have corresponding orthologs annotated within the genome of the strain for which the drug interaction pertains to. We then quantified the correlation between strain-specific drug impact and interaction scores for all 32 drug combinations, 3 of which were found to have a significant correlation (**Figure S4**, **Data S2**).

### PCA of strain- and media-specific prediction landscape

PCA was applied onto the strain- and media-specific drug interaction outcome prediction data (3655 interactions x 162 strain-media pairs). PCA results were then visualized onto the 2D space defined by the first two principal components (**Figure 4**). The cumulative pairwise distance between strains within the PCA-defined space was calculated for each of the nine media conditions to determine which growth environment yielded the largest strain-to-strain differences in outcome predictions (**Table S1**).

### Statistical Analysis

Three statistical tests were used for downstream model interpretation based on the set of sigma-delta features explaining 95% of model importance. First, the two-sample Student’s t-test with equal variance was used to determine sigma features that statistically differ in mean value between synergistic or antagonistic drug interaction outcomes. For delta features, the chi-square test with Yates’ correction was applied to determine any associations between synergistic or antagonistic drug interaction outcomes and drug- unique effects on a single gene. Finally, a hypergeometric test was performed to determine KEGG pathways that were significantly represented by genes associated with the top model features (those explaining 95% of the variance). This test was conducted for KEGG pathways annotated from both the *E. coli* MG1655 genome (organism code: eco) and the *M. tb* H37Rv genome (organism code: mtu). Of note, the number of pathways annotated for each genome served as the population size specified in the hypergeometric test. Importantly, p-values from all three statistical tests were adjusted using the Benjamini- Hochberg correction^90^ to minimize the false discovery rate.

## Acknowledgments

This work was supported by faculty start-up funds from the University of Michigan (UM), R01AI150826, R56AI150826 and R21AI144536 from National Institute of Allergy and Infectious Diseases, R35GM137795 from National Institute of General Medical Sciences, UM Precision Health, UM Office of the Vice Provost of Research, UM Endowment for Basic Sciences Accelerator Award, UM Research Scouts Award and UM MCUBED to

S.C. We thank members of the Chandrasekaran lab for feedback on the project.

## Author contributions

Conceptualization: SC Methodology: CHC, DCC, KS, SC

Investigation: CHC, DCC, MRS, NMR, MAO Visualization: CHC

Supervision: ADB, SC Writing—original draft: CHC, DCC, SC

Writing—review & editing: CHC, DCC, MRS, NMR, MAO, KS, ADB, SC

## Data and materials availability

All code used within this work is provided through the TACTIC GitHub repository (https://github.com/sriram-lab/TACTIC). Additional information on datasets and results are available in the supplementary materials.

